# Functional Deficits are Explained by Plantarflexor Remodeling Following Achilles Tendon Rupture Repair: Preliminary Findings

**DOI:** 10.1101/290833

**Authors:** Josh R. Baxter, Todd J. Hullfish, Wen Chao

## Abstract

Achilles tendon ruptures are painful injuries that often lead to long-term functional deficits. Despite the prevalence of these injuries, the mechanism responsible for limited function has not yet been established. Therefore, the purpose of this study was to present preliminary findings that support a hypothesis that skeletal muscle remodeling is the driving factor of poor outcomes in some patients. Biomechanical and ultrasonography assessments were performed on a patient that presented with poor functional outcomes 2.5 years after a surgically-repaired acute Achilles tendon rupture. Single-leg heel raise function was decreased by 70% in the affected limb while walking mechanics showed no deficits. Ultrasonography revealed that the affected limb had shorter, more pennated, and less thick medial gastrocnemius muscles compared to the unaffected limb. A simple computational model of a maximal-effort plantarflexion contraction was employed to test the implications of muscle remodeling on single-leg heel raise function. Subject-specific fascicle length and pennation measurements explained deficits in ankle work and power that strongly agreed with experimentally measured values using motion capture. These preliminary findings support the hypothesis that skeletal muscle goes extensively remodels in response to a ruptured tendon, which reduces the amount of work and power the joint can generate. This multidisciplinary framework of biomechanical, imaging, and computational modeling provides a unique platform for studying the complex interactions between structure and function in patients recovering from Achilles tendon injuries.

## Introduction

Achilles tendon ruptures often lead to long-term functional deficits [1–4] despite improved treatment and rehabilitation protocols [5–7]. Tendon elongation following injury is considered to be a predictor of functional deficits in a single-leg heel raise during the first year following surgical repair [3]. While tendon elongation is an important clinical indicator of patient function, the underlying musculoskeletal mechanisms that dictate plantarflexor function in patients treated for Achilles tendon ruptures remain unclear. Animal models suggest that acute changes to the musculo-tendinous structures of the foot and ankle elicit rapid remodeling of skeletal muscle to preserve joint function [8,9], but plantarflexor morphology changes in response to Achilles tendon ruptures and the effects on patient function have not been reported in humans.

The purpose of this study was to present preliminary findings utilizing a multidisciplinary framework for linking plantarflexor remodeling with functional deficits in patients who have suffered Achilles tendon ruptures. To accomplish this, we quantified plantarflexor morphology and patient function using an integrated ultrasonography and motion capture approach [10]. Further, we simulated a maximal-effort plantarflexion contraction using a computational model [11] to test the effects of muscle remodeling on plantarflexor function. Using this framework, we began to tested our hypothesis that long-term functional deficits are associated with decreased muscle fascicle length and increased pennation.

## Methodology

A 27-year old male (1.83 m and 84 kg) with a poor clinical outcome 2.5 years following an acute Achilles tendon rupture participated in this IRB approved study. The ruptured Achilles tendon on the right leg was surgically repaired by another provider using an open-reduction within 1 week of the initial injury. The patient described inability to participate in recreational activities due to loss of ankle strength and reported experiencing no pain. Poor clinical outcomes were confirmed using a clinical outcome score (ATRS score of 49/100), an evaluation by a fellowship-trained foot and ankle surgeon, and the inability to perform a single-leg heel raise [12].

Morphological measurements of the medial gastrocnemius were made on images acquired using ultrasonography (Telemed SmartUs, Vilnius, Lithuania) on the affected and unaffected sides. Muscle fascicle pennation, length, and muscle belly thickness (Fig. 1) were acquired using an 8 MHz linear ultrasound probe (Telemed LV8-5L60N-2, Vilnius, Lithuania) [13]. These measurements of medial gastrocnemius morphology were acquired with the patient positioned prone with an extended knee and ankle resting off the end of a treatment table. A single investigator identified the deep and superficial aponeuroses as well as a single fascicle using a custom-written script (MATLAB, The Mathworks, Natick, MA). Lines were manually fit to each of these structures and extrapolated off of the image until the fascicle intersected both deep and superficial aponeuroses to define fascicle length [14]. The angle between the deep aponeurosis and fascicle was calculated to define muscle pennation [15]. Muscle belly thickness was calculated as the distance between the superficial and deep aponeuroses in the mid-section of the image.

**Figure 1.**
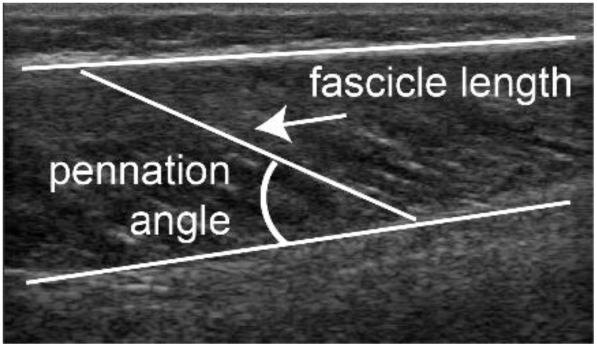
Ultrasound images of the medial gastrocnemius muscle were analyzed to quantify the fascicle length, pennation angle, and thickness (not shown for clarity).

Plantarflexor function was assessed through a battery of tests that consisted of isometric strength testing, walking, and single-leg heel raises. Isometric strength testing was performed in an isokinetic dynamometer (System 4, Biodex Medical Systems, Shirley, NY) with the patient seated with a fully extended knee and neutrally-aligned ankle [13]. Lower extremity biomechanics were quantified during comfortable speed walking and single-leg heel raises using a 12-camera motion capture system (Raptor series, Motion Analysis Corporation, Santa Rosa, CA) and 3 force plates (BP600900, Advanced Mechanical Technology, Inc., Watertown, MA). Fascicle shortening was synchronously acquired with an 8 MHz ultrasound probe during this battery of functional tests [16]. Plantarflexion kinematics, torque, power, and fascicle shortening dynamics were calculated to establish the link between muscular and patient function. Ankle biomechanics were calculated using open-source musculoskeletal software [17] (Opensim v3.3, Stanford University), and fascicle shortening dynamics were quantified using a custom-written tracking routine.

The effects of fascicle length and pennation on single-leg heel raise performance were tested using a simple computational model (Fig. 2A). To simulated a single-leg heel raise, maximal plantarflexion contractions of a foot pushing against a moving wall were simulated [11]. The wall retreated at a constant speed of 0.15 m/s, which was determined based on the average vertical velocity of the patient’s pelvis when performing single-leg heel raises on the unaffected limb. A single hill-type muscle model in series with an elastic element inserted into the foot and leg segments. Fascicle lengths and pennation angles were varied within the ranges of the patient’s affected and unaffected limbs (Table 1) to isolate the implications of these musculoskeletal parameters on plantarflexor function. Specifically, fascicle length and pennation angles were varied from 4 to 11 cm (1 cm increments) and 10 to 40 degrees (10 degree increments), respectively. Tendon slack lengths were determined geometrically by subtracting the medial gastrocnemius optimal fiber length from the muscle-tendon unit length when the ankle was positioned in 10 degrees plantarflexion [11]. Simulations began with the ankle neutrally aligned and completed when either the muscle stopped generating active force or when plantarflexion exceeded 50 degrees.

**Table 1.** Medial gastrocnemius morphology

|  | Unaffected | Affected | %change |
| --- | --- | --- | --- |
| Fascicle length | 11.2 cm | 4.6 cm | -59% |
| Pennation | 13° | 34° | 162% |
| Thickness | 2.9 cm | 2.2 cm | -24% |

**Figure 2.**
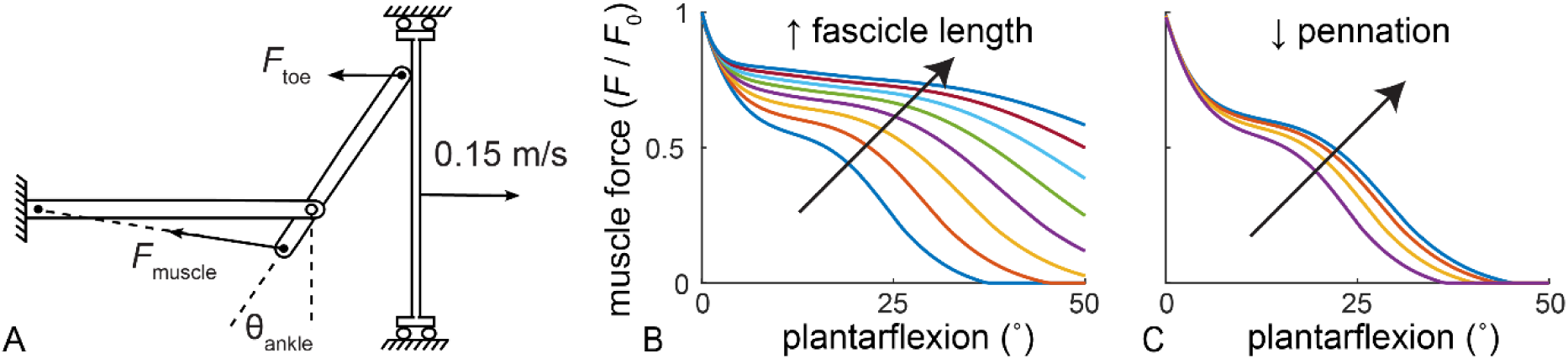
(A) A single-leg calf raise was simulated using a computational model of a foot maximally plantarflexing against a retreating wall. (B) Increasing fascicle lengths and (C) decreasing pennation increases the amount of muscle force that can be generated throughout the simulated single-leg calf raise. (B) pennation and (C) fascicle lengths were held constant at 10 degrees and 5 cm, respectively, to characterize the effects of changing a single muscular parameter. Muscle forces are presented as a ratio of the peak isometric force of the muscle (*F* / *F*_*0*_).

## Results

The affected limb demonstrated observable remodeling of the medial gastrocnemius muscle compared to the unaffected limb (Table 1). Medial gastrocnemius fascicle length decreased by 59%, pennation increased by 162%, and thickness decreased by 24% compared to the unaffected limb.

Plantarflexor function was compromised during isometric strength testing and single-leg heel raises but not during walking (Table 2). Isometric plantarflexor strength was 47% less in the affected side compared to the unaffected side. While the fascicles of the affected limb shortened 1.9 cm compared to 5.1 cm on the affected limb (63% decrease), relative fascicle shortening (41% – 46% resting lengths) was similar between sides (Table 2). Plantarflexion motion, torque, and power were similar between the affected and unaffected limbs during walking at 1.05 m/s. Medial gastrocnemius fascicles of each limb isometrically contracted (< 1 cm) throughout the stance phase. Severe deficiencies in plantarflexion motion, power, and muscle shortening explained a 70% decrease in single-leg heel raise height on the affected side (Table 2). The patient was only able to generate 12° of plantarflexion on the affected limb compared to 40° on his unaffected side. Despite no loss of peak plantarflexor torque, this 70% decrease in single-leg heel raise performance was explained by a 62% decrease in plantarflexor power. An 82% decrease of medial gastrocnemius fascicle shortening was measured using ultrasonography during the single-leg heel raise (Table 2).

**Table 2.** Plantarflexor biomechanics

|  |  | *Unaffected* | *Affected* | *%change* |
| --- | --- | --- | --- | --- |
| Isometric Strength | Torque (Nm) | 133 Nm | 70 Nm | -47% |
|  | MG shortening | 0.45 (5.1 cm) | 0.41 (1.9 cm) | ↔ (-63%) |
| Single Leg Heel Raise | Plantarflexion | 40° | 12° | -70% |
|  | Torque (Nm) | 140 Nm | 140 Nm | ↔ |
|  | Power (W) | 260 W | 100 W | -62% |
|  | MG shortening | 0.48 (5.4 cm) | 0.22 (1.0 cm) | -54% (-82%) |
| Walking (1.1 m/s) | Plantarflexion | 17 ± 1° | 18 ± 1° | ↔ |
|  | Torque (Nm) | 145 ± 6 Nm | 162 ± 5 Nm | ↔ |
|  | Power (W) | 284 ± 20 W | 313 ± 57 W | ↔ |
|  | MG shortening | < 1 cm | < 1 cm | ↔ |

Peak plantarflexor angle, torque, and power. Medial gastrocnemius (MG) relative shortening presented as ratio of absolute shortening / resting length and absolute shortening (in parentheses). Nominal % differences between the affected and unaffected sides are denoted by ↔.

Simulated single-leg heel raise function was found to be positively influenced by longer and less pennate fascicles (Fig. 3). Shorter and more pennate fascicles measured in the affected limb (Table 1) generated less force and power, which resulted in a 61% decrease in muscle work performed. When the 24% decrease in medial gastrocnemius thickness was factored into this analysis, the simulated affected limb did 70% less work than the unaffected limb, which strongly agreed with the 70% decrease in single-leg heel raise performance measured experimentally.

**Figure 3.**
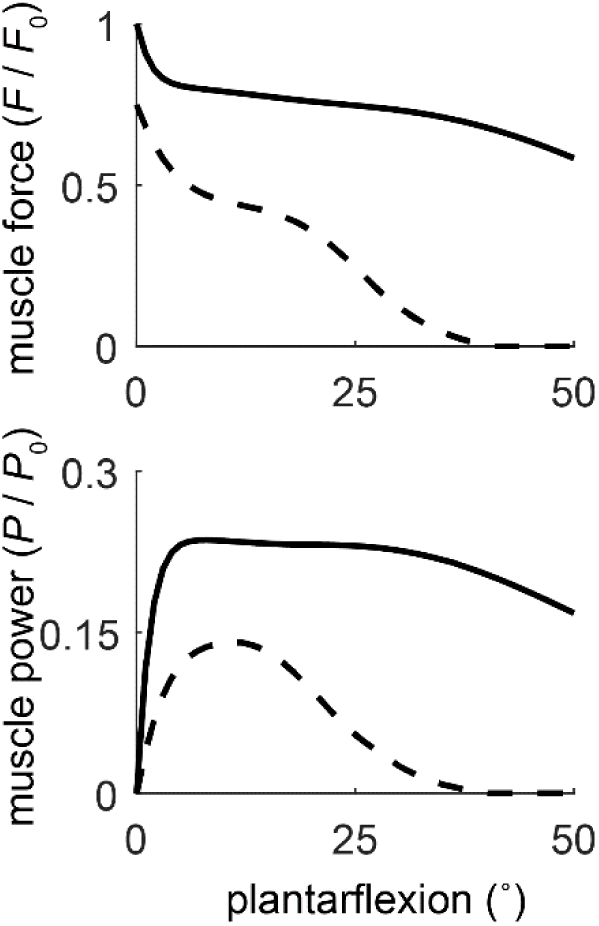
Patient-specific fascicle length, pennation, and muscle thickness values of the unaffected (solid line) and affected (dashed line) were tested in the simulation of a single-leg heel raise. Relative muscle force (*F* / *F*_*0*_, top) and power (*P*-*P*_*0*_, bottom) were greatly diminished with the shorter, more pennate, and less thick muscles experimentally measured in the patient’s affected limb. The amount of muscular work done by the affected limb decreased by 70% compared to the unaffected limb.

## Discussion

These preliminary findings support our hypothesis by demonstrating that morphological changes to the plantarflexors may explain long-term functional deficits in patients that suffer Achilles tendon ruptures. Clinical measures of function, such as the single-leg heel raise, directly test the plantarflexors ability to actively shorten and perform positive work [18,19]. On the other hand, cyclic activities, such as walking, can be successfully performed with reduced plantarflexor muscle function due to the well-described elastic energy return of the Achilles tendon [10,20]. Our experimental findings are supported by a computational model, demonstrating that plantarflexor morphology explains functional deficits when performing a single-leg heel raise.

Due to the noticeably smaller – and more proximal – gastrocnemius on the affected limb, we decided to acquire ultrasound images the same distance proximal of the muscle tendon junction on both limbs. This decision may explain the increased fascicle length on the unaffected limb. However, muscle pennation of the unaffected limb was within 1 standard deviation of reported values [13], suggesting that ultrasound images were acquired at an appropriate location. Further, results from our simple computational model strongly agree with experimental measures of patient function, further supporting these large disparities in fascicle lengths, pennation, and muscle thickness between the affected and unaffected side.

Traditional gait analysis may be a poor approach to assess patient function following an Achilles tendon rupture. We observed stereotypical plantarflexor function during gait [10] demonstrated that the patient walked efficiently despite severely compromised plantarflexors on the affected side.

Skeletal muscle is known to rapidly remodel in response to changes to surrounding musculoskeletal structures. Small animal models demonstrate that skeletal muscle responds to acute injuries by changing the number of sarcomeres in series in as little as one week [8]. Increasing the mechanical advantage of the tibialis anterior elicited morphological remodeling in order to maintain joint function [9]. However, these small animal models surgically altered the shortening demands by releasing the retinaculum, effectively increasing the tendon travel required for a given joint rotation. Unlike controlled animal experiments, tendon ruptures are traumatic injuries that elicit complex biologic responses [21] and compensatory strategies to unload the injured tissue [4]. Tendon ruptures are linked with long-term tendon elongation [3], which places unique shortening demands on the plantarflexors that appear to elicit detrimental morphological adaptations.

Poor long-term outcomes have been linked with tendon diastasis size at initial evaluation of injury [22]. This prior report suggests that smaller changes to the mechanical demands of the plantarflexors can be overcome while greater changes elicit detrimental adaptations to muscle morphology. Our findings suggest that the muscle remodels in order to maintain resting tendon tension, which leads to shorter and more pennate plantarflexor muscles – resulting in reduced functional capacity (Fig. 2). Minimizing tendon diastasis should be a priority, which can be modified based on surgical technique [23] and rehabilitation strategies [24,25]. Surgical interventions [26] and strength training [27–29] may restore fascicle morphology and improve patient function. Patients with impaired plantarflexor function may benefit from targeted strength training that elicits fascicle lengthening; however, the elongated tendon is a counter-acting constraint that appears to make longer muscle fascicles less feasible.

This study was affected by several limitations. These preliminary findings propose a mechanism for functional limitations in patients who suffered an acute Achilles tendon rupture explained by morphologic changes to the triceps surae. While this report has a small sample size (N = 1), the findings are supported by a computational model that closely quantifies ankle joint function based solely on changes to plantarflexor morphology. Morphological and functional measurements were not assessed at the time of injury because this patient was treated for the initial injury by another provider. It is possible that the unaffected limb underwent morphological changes in response to an increased workload to compensate for the functional constraints imposed by the injured limb; however, prior work does not support this theory [30]. Due to the noticeably smaller – and more proximal – gastrocnemius on the affected limb, we decided to acquire ultrasound images the same distance proximal of the muscle tendon junction on both limbs. This decision may explain the increased fascicle length on the unaffected limb. However, muscle pennation of the unaffected limb was within 1 standard deviation of reported values [13], suggesting that ultrasound images were acquired at an appropriate location. Further, results from our simple computational model strongly agree with experimental measures of patient function, further supporting these large disparities in fascicle lengths, pennation, and muscle thickness between the affected and unaffected side.

## Conclusion

Long-term functional deficits in patients who have suffered acute Achilles tendon ruptures may be explained by morphological changes in the plantarflexor muscles. We propose a multidisciplinary framework to rigorously quantify muscle remodeling in response to Achilles tendon injuries. This approach provides a means to prospectively study the implications of injury, treatment, and rehabilitation on plantarflexor function. Using this framework, we hypothesize that these structural changes are compensatory to tendon elongation. If correct, minimizing tendon elongation throughout treatment and rehabilitation is essential for long-term patient function. Linking plantarflexor structure and function in a prospective study is necessary to optimize treatment and rehabilitation options for patients with Achilles tendon injuries.

